# Glucose stockpile in the intestinal apical brush border in *C. elegans*

**DOI:** 10.1101/2023.08.11.553010

**Authors:** Takumi Saito, Kenji Kikuchi, Takuji Ishikawa

**Affiliations:** Graduate School of Biomedical Engineering, Tohoku University, Japan; Graduate School of Engineering, Department of Finemechanics, Tohoku University, Japan; Department of Molecular Biophysics and Biochemistry, Yale University, USA; Nanobiology Institute, Yale University, USA

**Keywords:** Diffusion coefficient, Glucose transport, Fluorescent glucose, Intestine, Fluorescence recovery after photobleaching, *Caenorhabditis elegans*

## Abstract

Since understanding the mechanisms of glucose transport is a crucial approach for pathological diseases induced by glucose toxicities such as diabetes, numerous studies have unveiled molecular functions involved in glucose transport in the nematode *Caenorhabditis elegans*, a commonly used model organism. However, physicochemical behaviors of glucose in intestinal lumen-to-cell are still elusive. To address that, we here evaluated a diffusion coefficient of glucose in the intestinal apical brush border in *C. elegans* by fluorescence recovery after photobleaching (FRAP) with fluorescent glucose. Our results indicate that the fluorescent glucose is accumulated in the intestinal brush border, where the diffusion coefficient of glucose is ∼10^−8^ cm^2^/s, two orders of magnitude slower than that in bulk. Namely, the intestinal brush border is considered to be a highly viscous layer composed of intestinal microvilli and more dominantly glycocalyx. These findings imply that this high concentration of glucose in the viscous layer acts as a stockpile.

## Introduction

The nematode *Caenorhabditis elegans* is the widely useful model organism due to its visibility and minimal construct with several organs and tissues comprised of 959 cells rather than other animals (Kaletta and Hengartner, 2006). Since the intestine in animals is one of the essential organs for nutrients transport and particularly absorption, the intestine in *C. elegans* has been used in many research fields such as molecular to multicellular biology, tissue developments, and disease pathogenesis and biomedicine, as is often related to human diabetes and glucose-related toxicity.

*C. elegans* under high glucose concentration indicated a reduction of life span (Kingsley et al., 2021; Schlotterer et al., 2009), representing mitochondrial dysfunction (Alcántar-Fernández et al., 2019). Regarding the molecular mechanism involved in the glucose transport, human GLUT2, one of the class 1 GLUT family and mainly localizes in the basolateral membrane of epithelial cells in human intestine and kidney, has been characterized as a passive transporter in a sodium-independent manner (Cheeseman, 1993; Thorens, 1992; Thorens and Mueckler, 2010). Likewise, human GLUT2-like glucose transporter FGT-1 has been identified in the intestinal basal membrane in *C. elegans* (Feng et al., 2013; Kitaoka et al., 2013). In addition, the human insulin/IGF-like signaling pathways have been reported to regulate FGT-1 expression in *C. elegans* (Kitaoka et al., 2016). Furthermore, the membrane-associated proteins, PAQR-2 and IGLR-2, have been recently reported to be the cooperative glucose sensitivity on the intestinal apical membrane in *C. elegans* (Devkota et al., 2021; Koyiloth and Gummadi, 2022; Svensk et al., 2016). Together, the glucose absorption of the intestine plays an important role in the physiological and pathological modulation of glucose levels. However, the dynamic behaviors of glucose in the intestinal lumen and cellular apical membrane are still unclear despite the first uptake to intestinal cells. While we have recently evaluated that the reciprocating flow in the intestinal lumen caused by the defecation motor program induces the nutrition uptake (Suzuki et al., 2022), how the glucose behaves in the microenvironment between the lumen and apical membrane known to be an intestinal brush border; and then how it is delivered to the cells with passive and/or active membrane-associated transporters are still open questions. To address them, we here carried out diffusion measurements of fluorescently labeled glucose in the intestinal brush border in adult worms by using fluorescence recovery after photobleaching.

## Material and methods

### *Caenorhabditis elegans* strains and culture methods

Wild-type strain N2 and mutants *erm1::GFP (mib15[erm-1::eGFP]) I, erm1-T544A::GFP (mib16[erm-1[T544A]::eGFP]) I*. and *erm1-T544D::GFP (mib16[erm-1[T544D]::eGFP]) I*. were all obtained from the *C. elegans* Genetics Center (CGC; University of Minnesota). In all cases, worms were maintained on the *E. coli* OP50-1 spread nematode growth media (NGM) plates at 20°C according to the typical protocol.

### Worms and sample preparation for imaging

Worms were collected from NGM plates to 1.5 mL flat tubes by using 700 µL of M9 buffer (Fig. 1A). 10 µL of M9 buffer including adult worms were transferred from the tube to a *ϕ*30 plastic dish. M9 buffer was then added up to 2 mL into the dish. Red fluorescent glucose (Glucose Uptake Assay Kit-Red; Dojindo Laboratories) was solved in 40 µL of dimethyl sulfoxide according to the manufacture’s protocol, and which was diluted with M9 in the dish (final concentration; 0.1 % (v/v)).

**Figure 1.**
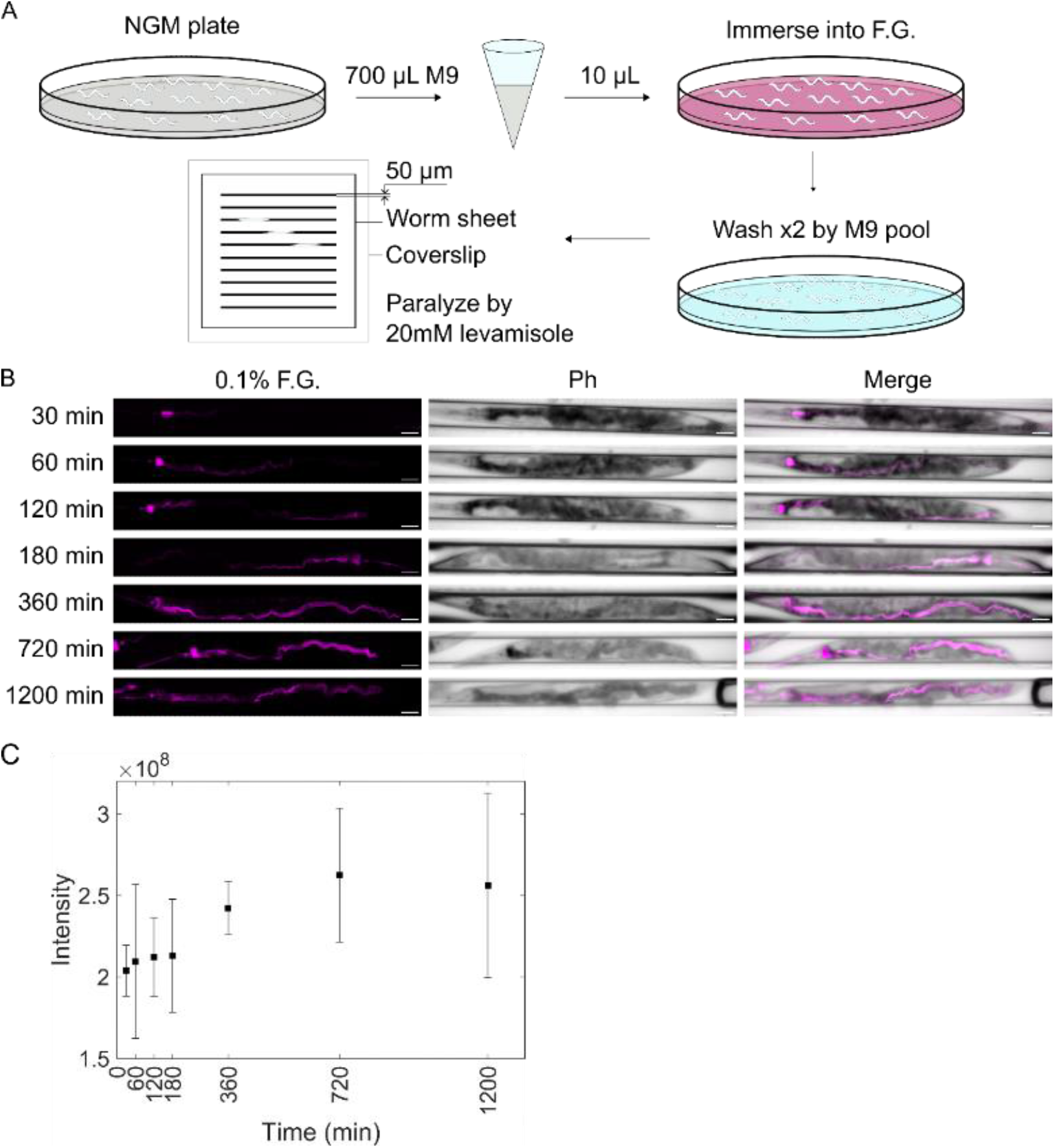
The red fluorescent glucose-uptake in the intestinal lumen of worms. (A) Briefly illustrating culture and preparation methods. Adult worms were cultured on typical NGM plates, collected and directly immersed in 0.1 % fluorescent glucose (F.G.) in M9 buffer, washed twice with M9 buffer, and paralyzed by 20 mM levamisole on the worm sheet for imaging. (B) Z-axis slice images of adult worms at different immersion times were acquired by using the confocal laser microscope and summarized along z-axis. Scale bars; 50 m. (C) The total intensity within worms at the different immersion times (n=4, 6, 5, 8, 6, 7 and 5 for 30, 60, 120, 180, 360, 720 and 1200 min, respectively).

These worms were incubated at 20 °C for a certain time (3-6 h for FRAP experiments) and then washed twice with M9 buffer and transferred onto PDMS worm sheet (Worm Sheet 50; Biocosm). Levamisole (20 mM in M9) drop was put on the sheet to paralyze the worms. Then, the sheet was gently covered with a coverslip for imaging (Fig. 1A).

We used PlasMem Bright Green (Dojindo Laboratories), a water-soluble plasma membrane dye, to visualize the apical membrane of intestinal lumen of worms. The ready-to-use PlasMem Bright Green was diluted to be 2% (v/v) with M9 in the dish immersing worms.

For FRAP experiments to estimate the diffusion coefficient of fluorescent glucose in bulk, fluorescent glucose solved in DMSO was diluted with glycerol (final concentration; 0.1% (v/v)). In addition, Rhodamine B (RhoB, 100 µM in DMSO) is diluted with glycerol (final concentration; 0.1 % (v/v)) to estimate the molecular weight of fluorescent glucose via diffusion measurements.

### Fluorescence images and FRAP measurements

Images were acquired by using a confocal laser scanning microscope (FV1000; Olympus) and 20x/NA0.75, 60x/NA1.00 or 100x/NA1.45 objectives to shoot two-dimensional fluorescent images for laser induced fluorescent (LIF) analysis (Kikuchi et al., 2019) and FRAP measurements (Saito and Deguchi, 2023). 488- and 561-nm wavelength lasers were used for both GFP and PlasMem Bright Green and for the red fluorescent glucose, respectively. In FRAP experiments, images were acquired for 5-8 s where the first two frames were pre-bleach images. A circular-shaped photobleaching was performed for 10-50 ms with both 405- and 488-nm wavelength lasers. The third frame became thus the post-bleach image, namely *t* = 0 . After FRAP experiments, worms are still survived as previously reported (Devkota and Pilon, 2018; Shivers et al., 2017; Walser et al., 2015).Unless otherwise the notice, images were analyzed with FIJI/ImageJ (NIH).

### FRAP analysis

Since the intensity profile often exhibits the Gaussian distribution in the circular-shaped photobleaching, we used the following equation:

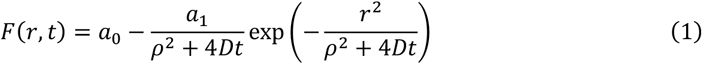

where *D* represents the diffusion coefficient, and *a*_0_, *a*_1_ and *ρ* are photobleaching parameters that describe the Gaussian distribution at *t* = 0 (Jonasson et al., 2008). These parameters were determined by fitting the spatiotemporal intensity distribution with a maximum likelihood method with MATLAB (MathWorks) (Deschout et al., 2010; Jonasson et al., 2008; Jonasson et al., 2010; Saito et al., 2021).

### Statistical analysis

Statistical differences were calculated by ANOVA, the Student’s *t*-test for variables with a Gaussian distribution, or the Mann-Whitney *U*-test for variables with a non-Gaussian.

## Results

### Direct immersion of worms in fluorescent glucose induces the glucose uptake

We have previously reported that worms fed *E. coli* OP50-1 labeled with the fluorescent glucose, visualizing glucose behaviors in the intestine of living worms (Suzuki et al., 2022). To test whether worms directly feed the fluorescent glucose without *E. coli*, here, the worms were directly immersed in the fluorescent glucose diluted in M9 buffer for 30, 60, 120, 180, 360, 720 and 1200 min at 20 °C (Fig. 1A). The worms were transferred onto the worm sheet, paralyzed with 20 mM levamisole, and observed by using the confocal laser scanning microscope equipped with the 20x objective lens. The z-slice images were acquired every 5 µm and total 90 µm in height (i.e., 18 slices), in which the intensity was summarized at every pixel along z-axis (Fig. 1B). The fluorescent glucose exhibited the uptake only in the pharyngeal at 30 min and then distributed in the intestine at the later cases. The total intensity of the fluorescent glucose in the whole body was indeed increased, indicating that worms fed the fluorescent glucose without *E. coli* (Fig. 1C). However, the slope of the uptake was decayed in 720-1200 min immersion (Fig. 1C). Therefore, 180-720 min immersed worms that are considered to actively feed the fluorescent glucose were used for the following experiments. In addition, the worms were immersed in the sodium fluorescein (uranine), resulting in that the uranine exhibited the high intensity in both the intestinal lumen and the intestinal cells (Fig. S1). In contrast to the uranine, the fluorescent glucose was dominantly accumulated in the intestinal lumen.

### Fluorescent glucose localizes in the intestinal apical brush border

The worms were immersed in both the fluorescent glucose and PlasMem Bright Green (Figs. 2A; S2). According to the correlation analysis, the fluorescent glucose was localized in the intestinal apical membrane (Fig. 2B). In addition, the mutants expressing ERM-1::GFP, the actin-crosslinking protein to facilitate the microvilli formation on the apical plasma membrane, was immersed in the fluorescent glucose, resulting in that the fluorescent glucose was localized in the microvilli (Fig. 2C,D). In both cases, the large correlation index between the fluorescent glucose and either PlasMem Bright Green or ERM-1::GFP was obtained (Fig. 2E). To test whether the fluorescent glucose was accumulated in or out of intestinal cells, the laser ablation was moreover performed onto the apical membrane. Consequently, the fluorescent glucose passively leaked out from the high concentration to the intestinal cells (Fig. S3). These results suggest that the fluorescent glucose was accumulated in the intestinal brush border where the apical membrane was penetrated by the ERM-1-anchored actin filaments.

**Figure 2.**
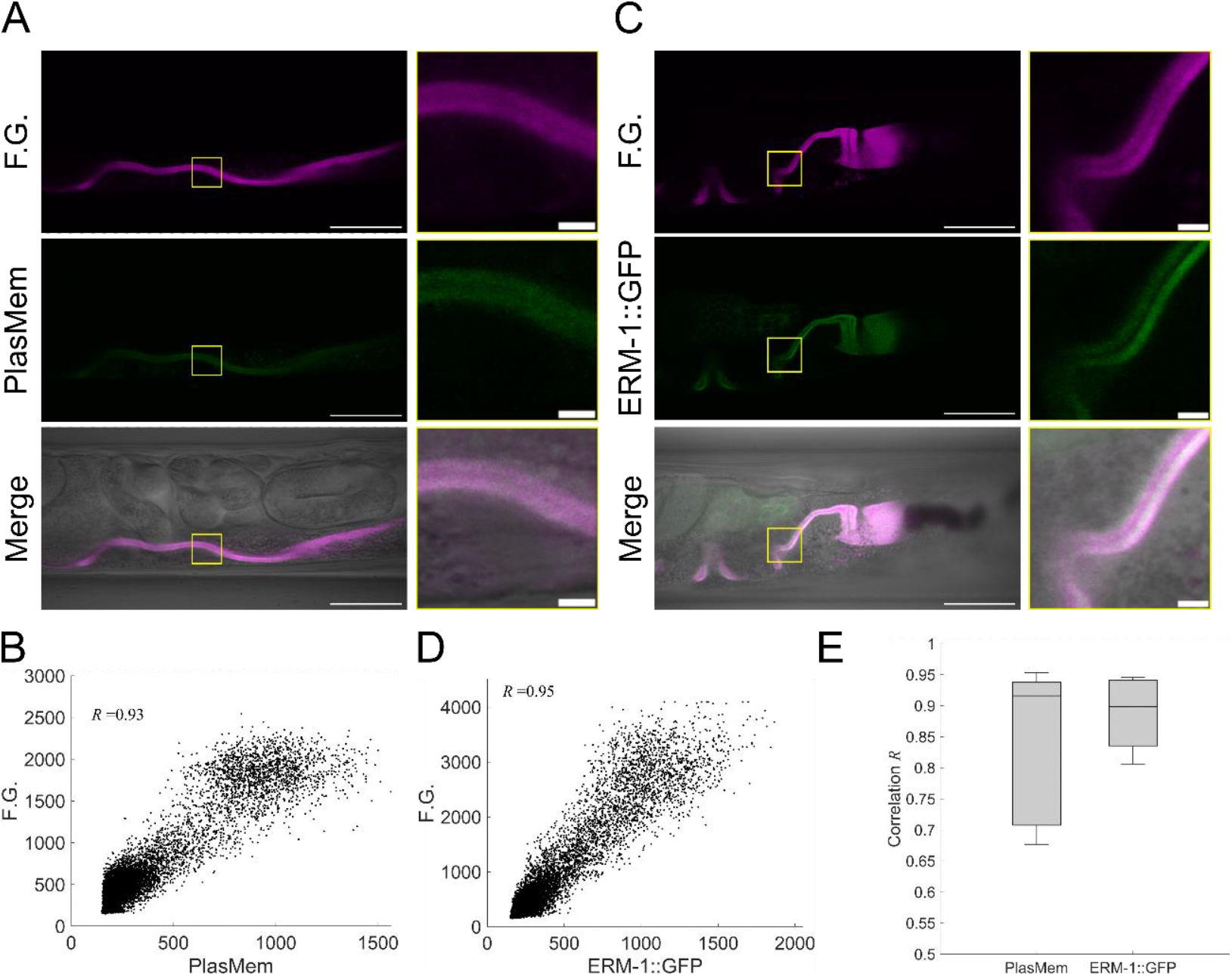
The fluorescent glucose (F.G.) was localized in the brush border on the intestinal apical membrane. A) Adult worms of wild type (WT) were immersed for 3 h in M9 buffer containing 0.1 % F.G. and 2 % PlasMem Bright-Green (PlasMem). Scale bars; 50 m (left) and 5 m (right). (B) Colocalization analysis between F.G. and PlasMem. (C) Adult worms of *erm-1::GFP* mutant were immersed for 3 h in M9 buffer containing 0.1 % F.G. Scale bars; 50 m (left) and 5 m (right). (D) Colocalization analysis between F.G. and ERM-1::GFP. (E) The correlation coefficient between the F.G. and either PlasMem or ERM-1::GFP (n=8 and 7 for WT worms immersed in PlasMem and *erm-1*::GFP mutants, respectively).

### Diffusion coefficient of glucose is two orders of magnitude slower than that in bulk

FRAP experiments were carried out onto the fluorescent glucose in the intestinal brush border in both the wild-type (WT) and the mutant (*erm-1::GFP*) adult worms (Fig. 3A). The spatiotemporal changes of the intensity were fitted by using Eq. (1) with the maximum likelihood method (Fig. 3B,C), determining the diffusion coefficient and other photobleaching parameters. The relatively slow diffusion coefficient of fluorescent glucose was observed in both worms, however, there is no significance between the WT and *erm-1-GFP* mutant worms, meaning that labeling GFP at *erm-1* does not affect the behavior of fluorescent glucose (Fig. 3D).

**Figure 3.**
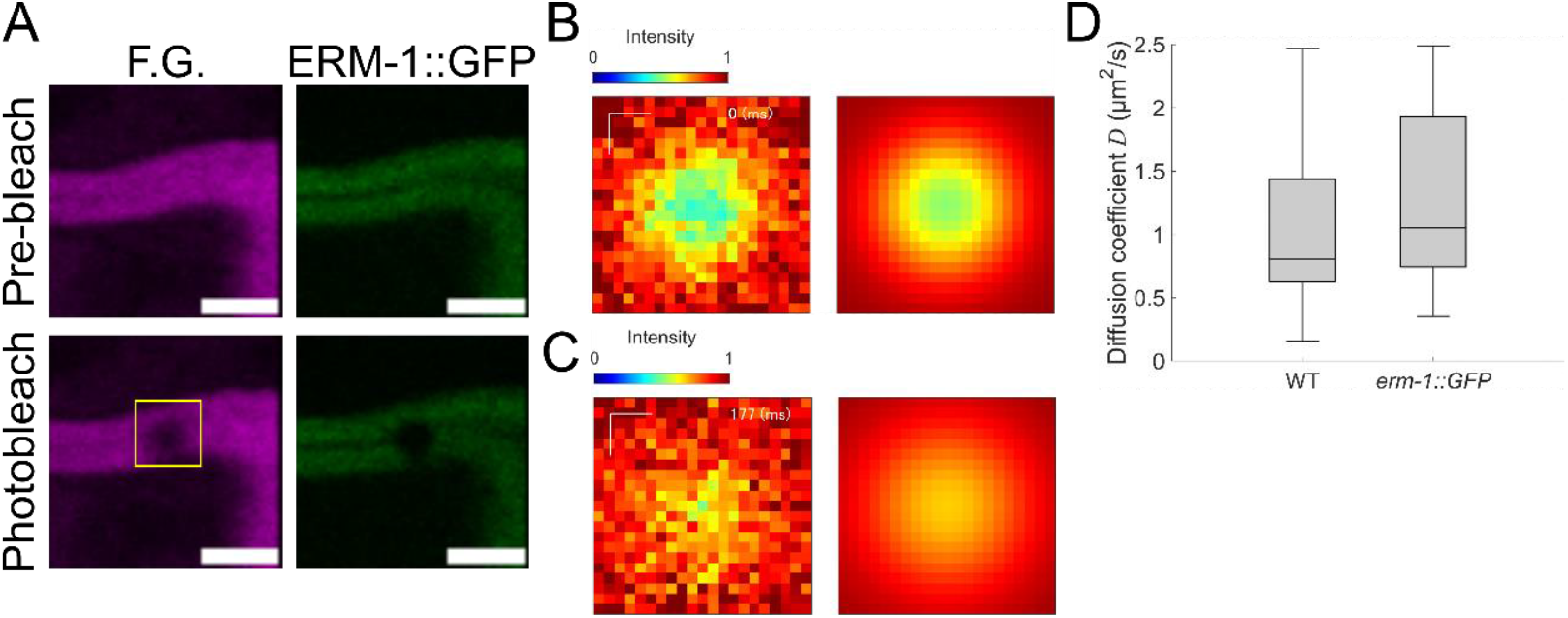
The diffusion coefficient of F.G. was determined by using FRAP experiments in the brush border on the intestinal apical membrane in wild type worms (WT) or *erm-1::GFP*. (A) Intensity distribution of F.G. (left) and ERM-1::GFP (right) in the mutants at pre-bleach (top) and photobleaching (bottom). (B and C) Intensity distribution in the yellow rectangles was normalized by the intensity if the pre-bleach frame (left), and analyzed by the diffusion equation (right) at given time points. Scale bars; 1 µm. (D) The diffusion coefficient of F.G. in WT and *erm-1::GFP*. There were no significant differences between WT and *erm-1::GFP* (*p* = 0.35 with *n*=34 and 7 for WT and *erm-1::GFP*, respectively).

Compared to the diffusion coefficient of glucose in water (*D* = 600 m^2^/s), the fluorescent glucose was approximately two-order of magnitude slowed in the intestinal brush border. Since the fluorescent glucose provided by the manufacture (Dojindo Laboratories) is considered to be comprised of at least two parts; a fluorophore and glucose, the molecular weight of fluorescent glucose must be larger than the “pure” glucose. Therefore, we first estimated this weight difference between the fluorescent glucose and glucose by measuring the changes in the diffusion coefficient. According to the Stokes-Einstein equation, the diffusion coefficient is in inverse proportion to the particle radius, and which is rewritten as 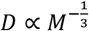 where *M* is the molecular weight. To estimate the molecular weight of fluorescent glucose, rhodamine B (RhoB) was thus used for the control molecule that represents known molecular weight (*M*_RhoB =_ 479.02 g/mol). FRAP experiments were then performed in the glycerol solution containing either 0.1 % (v/v) fluorescent glucose (Fig. 4A,B) or 0.1 % (v/v) RhoB (Fig. 4C,D). Consequently, the diffusion coefficient of both fluorescent glucose and RhoB was determined by fitting the spatiotemporal changes of the intensity to Eq. (1) with the maximum likelihood (Fig. 4E). By using the Stokes-Einstein relation and measured quantities, the molecular weight of fluorescent glucose was determined: *M*_FG =_ 636 ± 88.1 g/mol. Subtracting the molecular weight of glucose (180.15 g/mol), the subunit of fluorescent glucose was approximately 450 g/mol in weight, which was comparable to the molecular weight of red fluorophore rhodamine.

**Figure 4.**
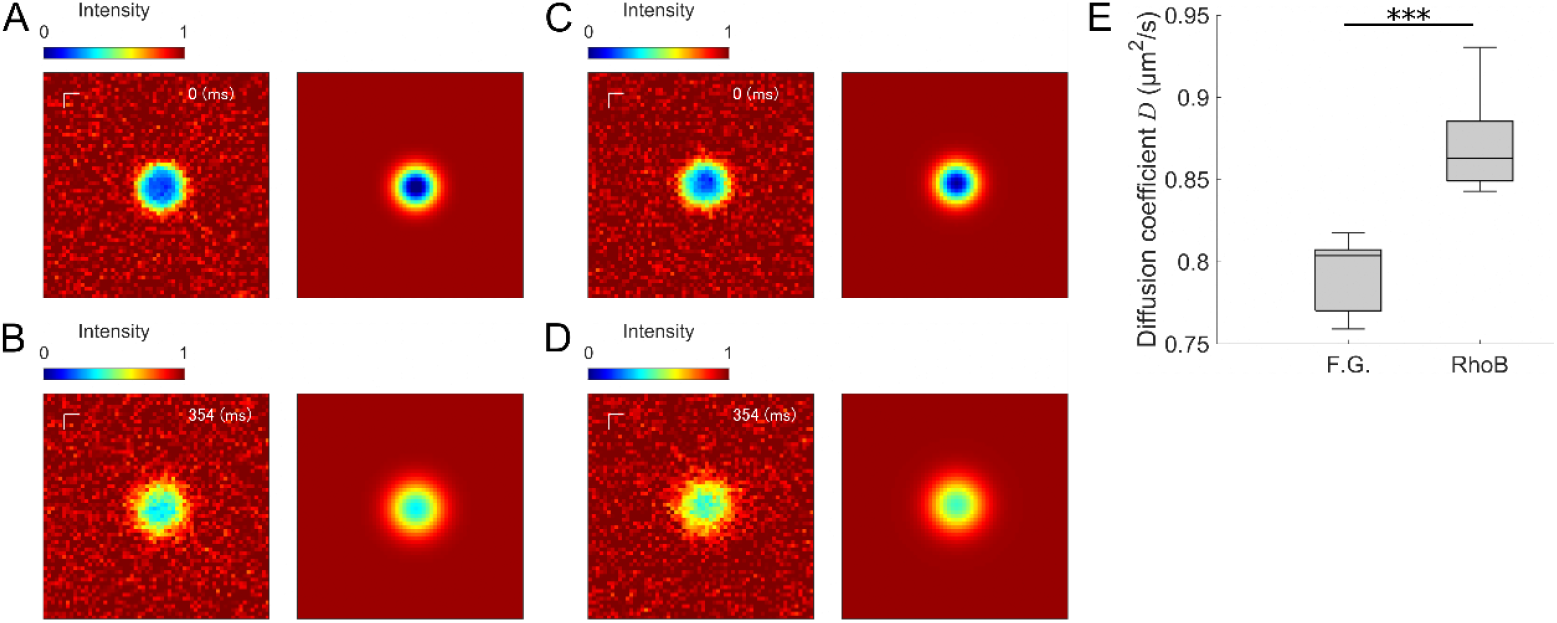
The diffusion coefficients of F.G. and Rhodamine B (RhoB) in the glycerol were determined by FRAP experiments. (A and B) Time changes of intensity distribution of F.G. normalized by the pre-bleach frame (left) and their regression distribution (right) at given time points; *t* = 0 (top) and *t* = 354 ms (bottom). Scale bars; 1 µm. (C and D) Time changes of intensity distribution of Rhodamine B (RhoB) normalized by the pre-bleach frame (left) and their regression distribution (right) at given time points; *t* = 0 (top) and *t* = 354 ms (bottom). Scale bars; 1 µm. (D) The diffusion coefficients of F.G. and RhoB in the glycerol, where *p* = 1.05 × 10^−4^ with *n* = 6 and 10 for F.G. and RhoB, respectively.

Secondly, we estimated the diffusion coefficient of fluorescent glucose in bulk. The Stokes-Einstein equation essentially allows to determine the changes in the diffusion under the different viscosity, namely *D* ∝ *η*^−1^ where *η* represents the viscosity. By using both the diffusion coefficient of fluorescent glucose measured in glycerol (Fig. 4E) and the viscosities of water *η*_w =_ 1.00 mPa.s and glycerol *η*_gly_∼900 mPa.s at 20 °C (Segur and Oderstar, 1951; Trejo González et al., 2011), the diffusion coefficient of fluorescent glucose in water was pragmatically estimated as approximately 700 µ m^2^/s that was comparable to the diffusion of pure glucose in water (Table 1). Altogether, the diffusion coefficient of fluorescent glucose in the intestinal brush border was two-order of magnitude slower than that in the bulk, meaning that the intestinal brush border was considered to be the highly viscous environment.

**Table 1.** The summary of quantities regarding the fluorescent glucose (F.G.).

| Diffusion coefficient of F.G. ($\mu\text{m}^2/\text{s}$) | | | Molecular weight (g/mol) |
| --- | --- | --- | --- |
| Brush border | Glycerol | Water |  |
| $1.02 \pm 0.64$ | $0.79 \pm 0.024$ | $\sim 700$ | $636 \pm 88.1$ |

### Mutation at C-terminal phosphorylation site of ERM-1 does not affect the diffusion of glucose

Next, we tested whether the changes in length of the intestinal microvilli affect the diffusion coefficient of fluorescent glucose. Since the point mutations at the C-terminal phosphorylation site; T544A and T544D as non-phosphorylatable and phosphomimetic variants, respectively, have been reported to shorten the length of the intestinal microvilli (Ramalho et al., 2020), the adult mutants of *erm-1[T544A]::GFP* and *erm-1[T544D]::GFP* were immersed in the fluorescent glucose. ERM-1[T544A]::GFP was colocalized with the fluorescent glucose (Fig. 5A,B), same as ERM-1::GFP does in the intestinal microvilli (Fig. 2). However, ERM-1[T544D]::GFP was expressed in not the intestine but the excretory canal with a weak GFP signal, while the fluorescent glucose exhibited the same localization pattern in the intestinal microvilli as shown above (Figs. 5C-E; Fig. S4). Interestingly, ERM-1[T544D]::GFP in the embryo and L4 larvae exhibited the localization in the intestinal apical membrane, which was consistent with the previously reported ERM-1 distribution (Ramalho et al., 2020) (Fig. S4). We performed FRAP experiments in both WT and the mutants, resulting in no significant differences among them in the glucose diffusion in the brush border (Fig. 5F).

**Figure 5.**
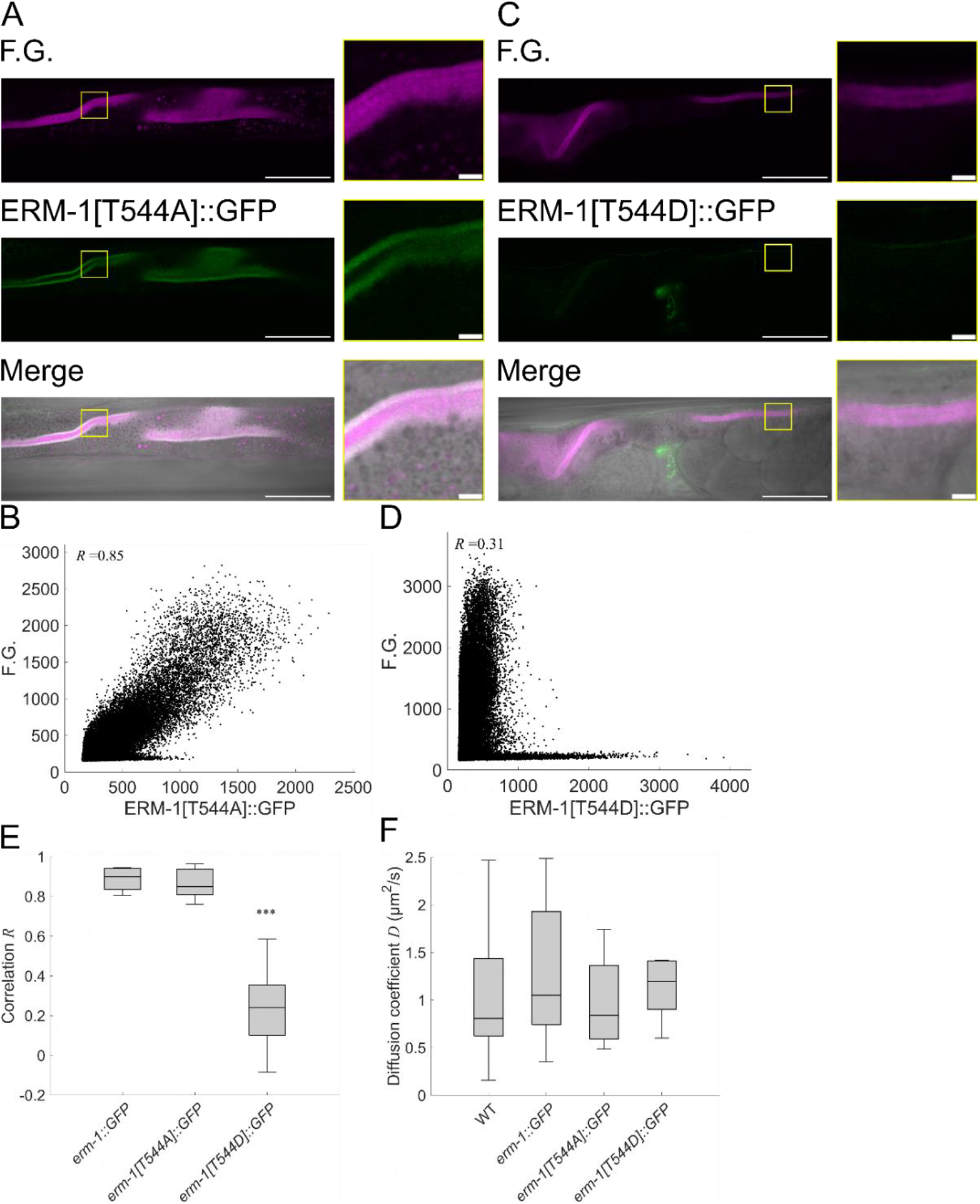
Colocalization analysis between F.G. and ERM-1::GFP in either non-phosphorylatable (*erm-1[T544A]::GFP*) or phosphomimetic (*erm-1[T544D]::GFP*) adult worms. (A) Fluorescence images of F.G. and ERM-1[T544A]::GFP and their merged images. Scale bars; 50 m (left) and 5 m (right). (B) Correlation analysis between F.G. and ERM-1[T544A]::GFP. (C) Fluorescence images of F.G. and ERM-1[T544D]::GFP and their merged images. Scale bars; 50 m (left) and 5 m (right). (D) Correlation analysis between F.G. and erm-1[T544D]::GFP. (E) Th correlation coefficient was significantly decreased in the phosphomimetic mutants. *p* < 10^−6^ (ANOVA) with *n* = 7, 5 and 10 for erm-1::GFP, erm-1[T544A]::GFP and erm-1[T544D]::GFP, respectively. (F) There were no significant differences (ANOVA) in the diffusion coefficient of F.G. on the intestinal brush border among the mutants (n=34, 7, 4 and 6 for WT, *erm-1::GFP, erm-1[T544A]::GFP* and *erm-1[T544D]::GFP*, reprectively).

## Discussion

To evaluate the glucose behaviors in the intestinal microenvironment in *C. elegans*, we performed the diffusion measurements with the fluorescent glucose by fluorescence recovery after photobleaching (FRAP). First, we observed the localization of the fluorescent glucose in the adult worms (Fig. 1). At the early stage of feeding, the fluorescent glucose existed in the pharyngeal (Fig. 1B), as is observed in the use of fluorescently labeled *E. coli* (Suzuki et al., 2022). Then, the worms showed the increase of the intensity in their intestinal lumen as we expected, meaning the *E. coli* independent glucose feeding (Fig. 1B,C). In addition, high concentration of the fluorescent glucose was observed in the intestinal lumen rather than the intestinal cells, unlike the localization of uranine (Fig. S1). These results presumably suggest the presence of active transporters that enable to control the amount of glucose uptake into the intestinal cells and/or the dysfunction of fluorophore due to its conformational changes by the consumption of glucose comprising the fluorescent glucose in the cells.

The multicolor imaging with the fluorescent glucose and either the lipophilic dye (PlasMem Bright Green) or ERM-1::GFP that anchors actin filaments at the apical plasma membrane revealed the localization of the fluorescent glucose in the intestinal microvilli (Fig. 2). Considering the crowded structures on the intestinal apical surface where the numerous microvilli are aligned every tens of nanometer scale (Bidaud-Meynard et al., 2021), it is still difficult to clarify whether the fluorescent glucose is accumulated around the luminal surface or inside the intestinal cells. Thus, we performed the laser ablation, resulting in that the fluorescent glucose was leaked from the apical region to the intestinal cells (Fig. S3). These results revealed that the fluorescent glucose is accumulated in the brush border on the apical surface. Indeed, this belt-shaped-like accumulation of fluorescent glucose was approximately ∼1 µm in width, which is consistent with the typical width of the brush border in worms (Khan et al., 2013; Ramalho et al., 2020). Therefore, the fluorescent glucose is considered to be trapped by a high viscosity layer in the brush border known to be glycocalyx including digestive and protective molecules (McGhee, 2007; Stutz et al., 2015; Zhou et al., 2023).

Indeed, FRAP experiments indicated that the diffusion coefficient of fluorescent glucose was two-order of magnitude slower than that in water (Fig. 3). To compensate the differences of diffusion coefficient between the fluorescent glucose and the simple glucose caused by the changes in their molecular weight, we also carried out FRAP in the glycerol. First, the diffusion coefficients of fluorescent glucose and rhodamine B (*M*_RhoB =_ 479.02 g/mol) were measured in the glycerol solution and hence the molecular weight of fluorescent glucose was determined by using the Stokes-Einstein equation. As we mentioned, the estimated molecular weight of fluorescent glucose was approximately 636 g/mol. By subtracting the molecular weight of glucose (180.15 g/mol), the change in the molecular size of fluorescent glucose is 450 g/mol by labeling a fluorophore. This estimation is reasonable because the molecular weight of representative red fluorophore rhodamine(s) is more or less the same as this weight difference. Moreover, the Stokes-Einstein equation estimated the diffusion coefficient of fluorescent glucose in water as approximately 700 µ m^2^/s by considering the differences of the viscosity (water: *η*_w_ =1.00 mPa.s and glycerol: *η*_gly_∼900 mPa.s at 20 °C (Segur and Oderstar, 1951; Trejo González et al., 2011)), which is obviously slower than that in water. Even though the molecular weight is three times larger than the glucose, the diffusivity is not obviously affected according to the Stokes-Einstein equation (i.e., 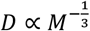). This high viscosity layer is considered to decrease the diffusion coefficient and then accumulate the glucose regardless of the fluorescence labeling. To test whether the microvilli in this high viscosity layer directly obstruct the diffusivity of glucose, we measured the diffusion coefficient in the mutants that express shorter microvilli than WT in length (Ramalho et al., 2020). Consequently, there are no significant differences between WT and the mutants in the diffusion coefficient (Fig. 5), meaning that the changes of microvilli in length do not have significantly much contribution to the diffusivity of glucose. In contrast, this result implies that the glycocalyx, the representative nanoscale molecular complex on the apical surface, predominantly decreases the glucose diffusion. However, the interplay between the glucose diffusion and functional molecular complexes including digestive and antiviral molecules in the glycocalyx is still unknown (McGhee, 2007).

Regarding the molecular mechanism in glucose transports, GLUT1-4 among the human GLUT family, have been characterized and well-studied as major passive transporters in the plasma membrane of several cells (Holman, 2020; Thorens and Mueckler, 2010). Especially, GLUT2 localizes in the basolateral membrane of epithelial cells in the intestine, where that delivers the glucose and fructose with the sodium-independent manner (Cheeseman, 1993). In contrast, sodium-dependent co-transporters SGLTs have been identified as active transporters, in which particularly SGLT1 is expressed in the intestinal apical membrane (Deng and Yan, 2016; Sano et al., 2020). In *C. elegans*, only the human GLUT2-like passive glucose transporter, FGT-1, has been identified as the homolog of GLUTs in the intestinal basal membrane (Feng et al., 2013; Kitaoka et al., 2013). In addition, the human insulin/IGF-like signaling pathways have been reported to regulate FGT-1 expression in *C. elegans* (Kitaoka et al., 2016). Together, such glucose transporters play an important role in the physiological and pathological modulation of glucose levels with both passive and active transports from cells to whole the body and from the intestinal luminal tube to cells, respectively. Besides that, numerous theoretical models have been provided for glucose passive and active transports (Naftalin and De Felice, 2012; O., 1966). For SGLT-like cotransporters, the cotransport models have been developed: SGLT1/Na+ cotransporter model (Eskandari et al., 2005; Mackenzie et al., 1998), the conventional antiporter models such as Na+:H+ exchange (Olkhova et al., 2006), and the network thermodynamic model (Mikulecky, 1983; Mikulecky, 2001). Particularly for SWEET1 transporter, the three state model was developed (Park et al., 2022), where the glucose concentration in the outside of membrane indicates the increase of glucose influx. The defecation motor program, which enables mechanical transport and mixing in the lumen, allows for providing intestinal contents along to the intestine and enhancing glucose uptake efficiently (Suzuki et al., 2022). Taking into account the insight from these models, our result ―the fluorescent glucose is accumulated in the viscous brush border― suggests that high glucose concentration allow to increase the stochastic glucose/transporters interaction, ending up high glucose influx to cells. However, carriers such as the passive and active glucose transporters in *C. elegans* intestine have not been identified. Thus, we here represent a simplest minimal model:

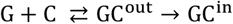

where *G* and *C* represent the glucose and carrier, GC^out^ and GC^in^ represent the glucose/carrier complexes in the inside and outside of cells. This model reduces the following equations:

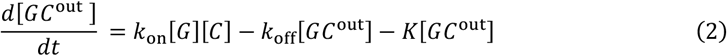

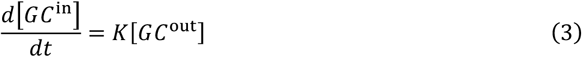

where *k*_on_ and *k*_off_ are stochastic on- and off-rates, and *K* is a constant value. This model again represents that high concentration of glucose in the outside of cells namely the brush border facilitates the glucose influx likewise the previously developed models (O., 1966; Park et al., 2022). To understand further details in the intestinal glucose transports in *C. elegans*, although FRAP unveiled macroscopic glucose behaviors in the brush border, genetically identifying and characterizing the glucose transporters are still open challenges.

## Acknowledgements

The authors thank Dr. Keiko Numayama-Tsuruta and Dr. Toshihiro Omori for their professional discussions. T. S. is supported by Japan Society for the Promotion of Science (JSPS).

## Autor contributions

Conceptualization: T. S., K. K., T. I.; Methodology: T. S., K. K.; Validation: T. S., K. K., T. I.; Formal analysis: T. S., K. K.; Investigation: T. S., K. K., T. I.; Resources: K. K., T. I.; Data curation: T. S., K. K.; Writing - original draft: T. S.; Writing – review & editing: T. S., K. K., T. I.; Visualization: T. S., K. K., T. I.; Project administration: K. K., T. I.; Funding acquisition: T. S., K. K., T. I.

